# Deconstruction of *Saccharomyces* hybrids reveals multiple genomic regions required for thermotolerance

**DOI:** 10.64898/2026.09.29.755322

**Authors:** Nilima Walunjkar, James H. Miller, Lydia Levesque, Justin C. Fay

## Abstract

The genetic basis of phenotypic differences between species is fundamental to our view of evolution. In contrast to variation within species, phenotypic differences between species can result from a long-term sequential process whereby prior steps and multiple changes at a single locus can play important roles. However, investigating the genetic basis of phenotypes that have evolved over long time periods is limited by reproductive barriers between species. In this study, we use hybrid genetic analysis to examine reproductively isolated *Saccharomyces* species that have diverged in their thermal growth limits over the last 15 million years. By selectively eliminating *S. cerevisiae* chromosomes in hybrids with *S. uvarum* or *S. kudriavzevii*, we show that most chromosomes cause a substantial loss of thermotolerance. Reciprocal loss of chromosome 5 (Chr 5) confirms that the hybrid’s thermotolerance depends on the *S. cerevisiae* chromosome and shows that it is partially inhibited by the *S. uvarum* chromosome. Using CRISPR-Cas9 to induce targeted loss of heterozygosity, we find that loss of either the left or right arm of Chr 5 results in loss of thermotolerance. Fine-mapping, complementation and reciprocal hemizygosity analysis identify *MMS21*, which encodes an essential subunit of the Smc5-Smc6 complex, as responsible for loss of thermotolerance associated with the left arm of Chr 5. Our findings suggest that divergence in thermotolerance over long time-periods is caused by multiple loci across the genome, with each playing a critical role in enabling growth at high temperatures.

**Author Summary:** How new traits evolve and give rise to differences between species is a central question in biology. Although many studies have identified genes that contribute to variation within species, much less is known about the genetic changes responsible for differences that accumulate between species over millions of years. One challenge is that species often become reproductively isolated, making it difficult to map and identify genes that contribute to evolved traits.

In this study, we used hybrids between different yeast species to investigate the evolution of thermotolerance, the ability to grow at high temperatures. By selectively removing chromosomes inherited from one species in the hybrid, we found that thermotolerance depends on many regions distributed across the genome. Focusing on one chromosome, we identified a gene called *MMS21* that contributes to growth at elevated temperatures, while also showing that additional regions on the same chromosome are required for thermotolerance.

Our results suggest that thermotolerance evolved through the accumulation of many genetic changes, each playing an important role in enabling growth at high temperatures. More broadly, our work demonstrates how hybrid genetic approaches can overcome barriers between species and provide insight into the genetic basis of traits that have evolved over long periods of evolutionary time.

## Introduction

The genetic basis of phenotypic differences between species is important to our understanding of evolution. The genetic changes underlying trait variation shape our view of whether evolution proceeds gradually through numerous changes of small effect or a few large effect changes [1,2]. They also provide insight into whether evolution is constrained to certain loci or evolutionary paths [3,4], and how often it is deterministic and repeatable [5,6]. However, studies have revealed a diverse range of genetic architectures underlying phenotypic divergence, making it important to delineate various evolutionary factors that influence the genetic basis of evolutionary change [7,8].

One important factor is the timescale over which phenotypic divergence occurs. While many patterns observed within or between closely related species may extend to distantly related species, important distinctions have emerged with respect to the type of traits that differ and their genetic basis. Phenotypic differences between sister species often involve rapidly evolving reproductive characters [9,10]. Moreover, genetic studies frequently examine coloration differences and find single genes of large effect [11–14]. Thus, our view of phenotype divergence is substantially influenced by interspecific studies of readily evolvable traits, such as those caused by loss of function mutations or introgressions of divergent alleles from other species [6]. Traits that evolved over longer evolutionary timescales warrant special attention as they may be shaped by the accumulation of multiple genetic changes and may evolve through mechanisms not easily accessed over shorter evolutionary periods.

A prominent result that has emerged from studies of divergence between species is that certain genes can have large phenotypic effects due the cumulative impact of multiple substitutions of small effect [6]. Multiple changes can be concentrated within a single gene when few genes influence a trait [15,16] or when a gene is a key modulator of regulatory networks with the right level of pleiotropy [8,17]. As large effect genes are easier to detect, it is difficult to disentangle the frequency of such genes evolving under a micromutational model and our ability to find them [1,6].

Reproductive isolation is a major barrier to genetic analysis of phenotypic divergence between species. While many closely related species are interfertile, genetic incompatibilities accumulate rapidly over longer periods [18]. For distantly related species that form viable but sterile hybrids, genetic analysis is often limited to reciprocal hemizygosity or deletion mapping [19–21]. One way to circumvent hybrid sterility is to use CRISPR-Cas9 to generate mitotic recombination within hybrids [22,23]. Transgenic approaches also avoid hybrid sterility but require candidate genes unless many genes can be screened or large genomic segments can be transferred [24]. Although these approaches have some limitations, they enable reproductive barriers to be overcome for genetic analysis in distantly related species.

The *Saccharomyces* yeast species provide an attractive system to dissect the genetic basis of interspecific phenotypic divergence and conspicuously differ in their thermal growth profiles. *S. cerevisiae*, the most thermotolerant species in this group can grow at temperatures ∼8 °C higher than two cryotolerant species *S. uvarum* and *S. kudriavzevii* [25–27]. Hybrids of these species are viable but sterile due to high nucleotide sequence divergence that inhibits recombination and incompatibilities between nuclear and mitochondrial genes [28]. However, hybrids are able to grow at high (37 °C) temperatures [29], making it possible to deconstruct thermotolerance by loss of *S. cerevisiae* chromosomes or chromosomal regions.

In this study we use hybrid genetic analysis to study the genetic basis of thermal divergence that has occurred over the last 15 million years in the *Saccharomyces* species. We induced loss of whole chromosomes in hybrids, thereby measuring the effect of entire *S. cerevisiae* chromosomes on growth at high temperature. We selected one chromosome of large effect and used CRISPR-based mitotic recombination to refine thermotolerance to regions on both the left and right arm of the chromosome. Plasmid complementation and reciprocal hemizygosity analysis identified *MMS21* as a major effect gene on chromosome 5. Our results do not exclude many genes of small effect, but show how hybrid thermotolerance requires the simultaneous presence of multiple chromosomal regions of large effect.

## Results

### Multiple *S. cerevisiae* chromosomes are required for hybrid thermotolerance

To map the genetic basis of thermotolerance, we generated a panel of *S. cerevisiae* x *S. uvarum* hybrids each missing a single *S. cerevisiae* chromosome. Loss of the target *S. cerevisiae* chromosome was generated by selecting against a *URA3* marker placed adjacent to each chromosome’s centromere [30](Figure 1). We isolated three to five replicate chromosome loss strains for twelve out of the sixteen chromosomes. From whole genome sequencing we found that in some strains loss of the target chromosome was accompanied by duplication of the homologous *S. uvarum* chromosome (n = 21) or aneuploidy of non-target chromosomes (n = 12, Table S1). There were two chromosomes (4 and 12) for which loss was always accompanied by a non-target aneuploidy. Loss of *S. cerevisiae* Chr 4 was always accompanied by duplication of *S. uvarum* chromosomes 2 and 4, and can be explained by a reciprocal translocation between Chr 2 and Chr 4 between the two species [31]. Loss of *S. cerevisiae* Chr 12 was always accompanied by duplication of *S. uvarum* Chr 10, although there are no translocations between these chromosomes.

**Figure 1:**
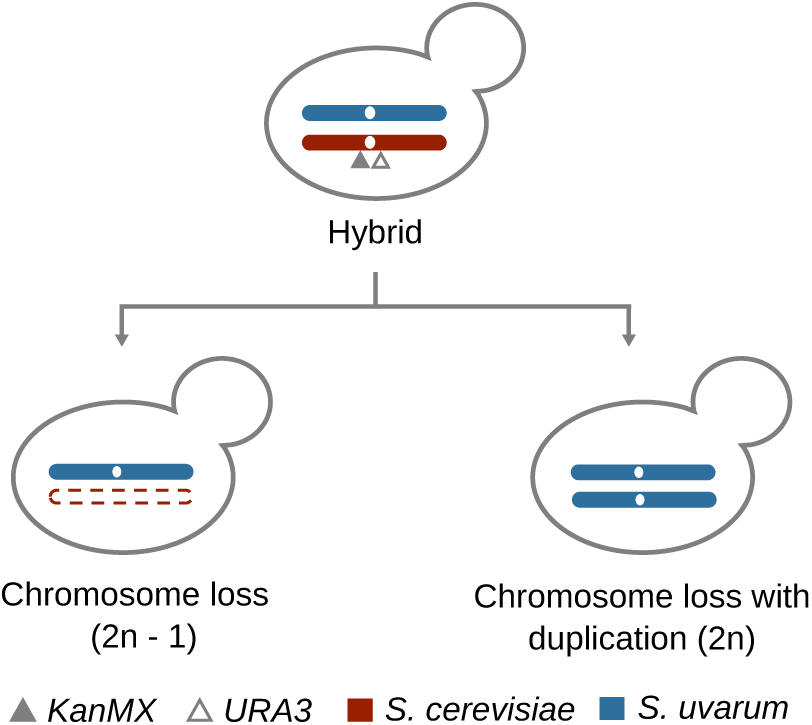
Targeted loss of *S. cerevisiae* chromosomes. Selective elimination of an *S. cerevisiae* chromosome from *S. cerevisiae* x *S. uvarum* hybrids with KanMX and URA3 at the centromere (open circle) results in loss of the target chromosome and 2n - 1 ploidy, or loss accompanied by duplication of the homologous *S. uvarum* chromosome, restoring ploidy to 2n.

We tested whether loss of *S. cerevisiae* chromosomes was associated with reduced thermotolerance. Growth was measured by colony sizes at a temperature (37 °C) above the thermal limit of *S. uvarum* and at a permissible temperature for both species (30 °C). To account for phenotypic variance of the hybrid parents (Figure 2A), all chromosome loss strains were normalized to their euploid hybrid parent. On dextrose (YPD), loss of eight of the twelve chromosomes tested caused a significant reduction in growth at high compared to the permissive temperature (p < 0.05, Figure 2B, Table S2).

**Figure 2:**
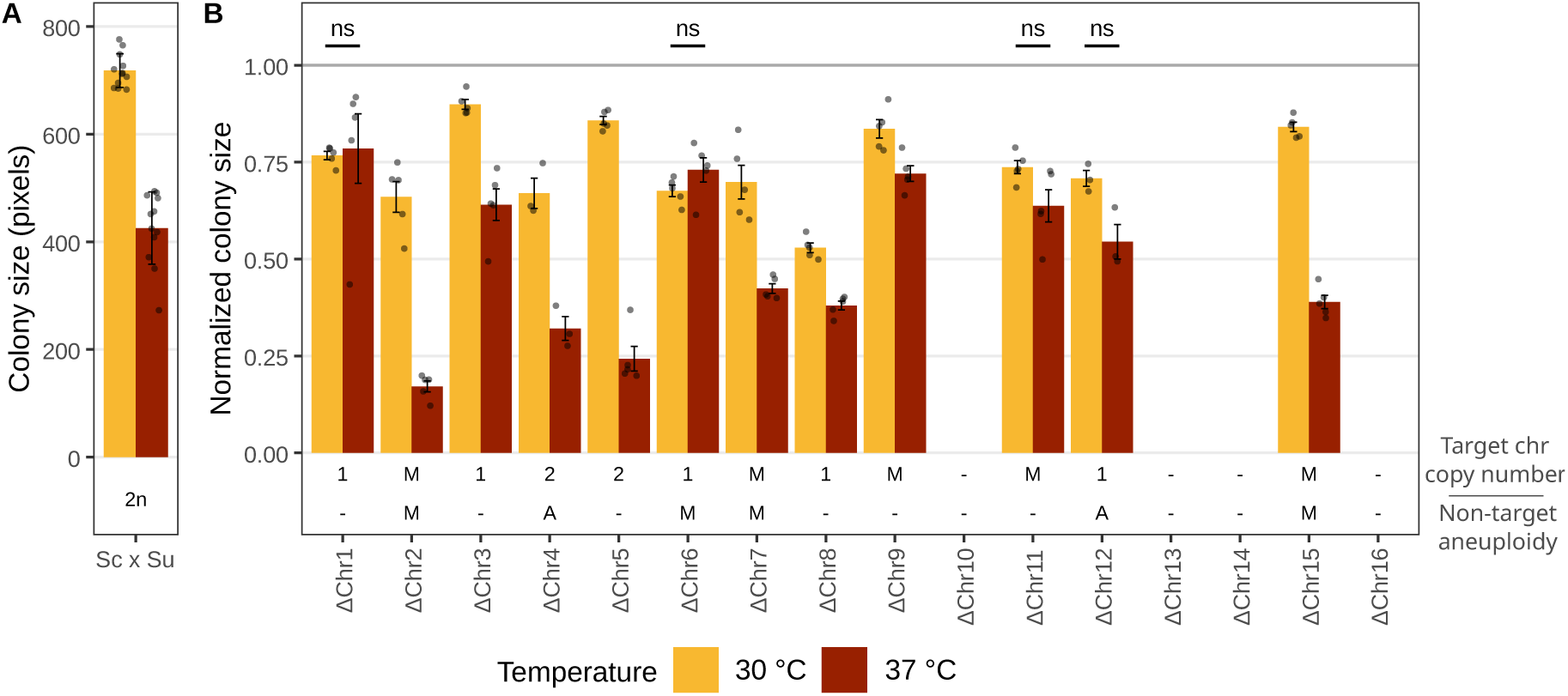
Loss of thermotolerance in *S. cerevisiae* x *S. uvarum* chromosome loss strains. Thermotolerance was measured by colony size after three days of growth on dextrose (YPD) at 37 °C versus 30 °C with bars representing the mean of replicates (grey points). (A) Colony size of euploid parental hybrids with error bars representing standard deviation. (B) Colony size of chromosome loss strains normalized to their euploid parental hybrid. Each targeted *S. cerevisiae* chromosome (x axis) has a significant effect on thermotolerance unless annotated as ns (Table S2). Error bars represent standard error of the mean. Annotations below the plot indicate the copy number of the target chromosome (1, 2, or M for mixed) and the presence of non-target aneuploidy (A = present, M = mixed).

At the permissive temperature, all chromosome loss strains showed a growth defect relative to their hybrid parents (Figure 2B). This growth defect could be caused by dosage differences in the target chromosome, non-target aneuploidy or the absence of *URA3*. We tested for an association with target chromosome dosage and found no growth difference between strains that had duplications of the *S. uvarum* homolog and strains with just a single copy of the target chromosome (p > 0.05, Table S3). Non-target aneuploidy also did not explain the growth defects in the chromosome loss panel (p > 0.05, Table S3). Thus, the lack of *URA3* likely explains growth defects at 30 °C for most of the chromosome loss strains.

We also measured thermotolerance on glycerol (YPG) which is assimilated by respiration. Similar to growth on dextrose, nine of the twelve chromosomes showed reduced growth at high temperature (p < 0.05, Table S2, Figure S1). However, on glycerol we found non-target aneuploidy (n = 12) is associated with reduced colony size at 30 °C (Table S3). To compare thermotolerance across substrates we measured thermotolerance as the normalized colony size at 37 °C relative to 30 °C. Only two of the tested chromosomes (1 and 11) had no significant effect on either substrate, and two chromosomes (6 and 12) had no effect on dextrose but a large effect on glycerol. Overall, loss of a *S. cerevisiae* chromosome had a larger effect on high temperature growth on glycerol compared to dextrose (Figure 3A).

**Figure 3:**
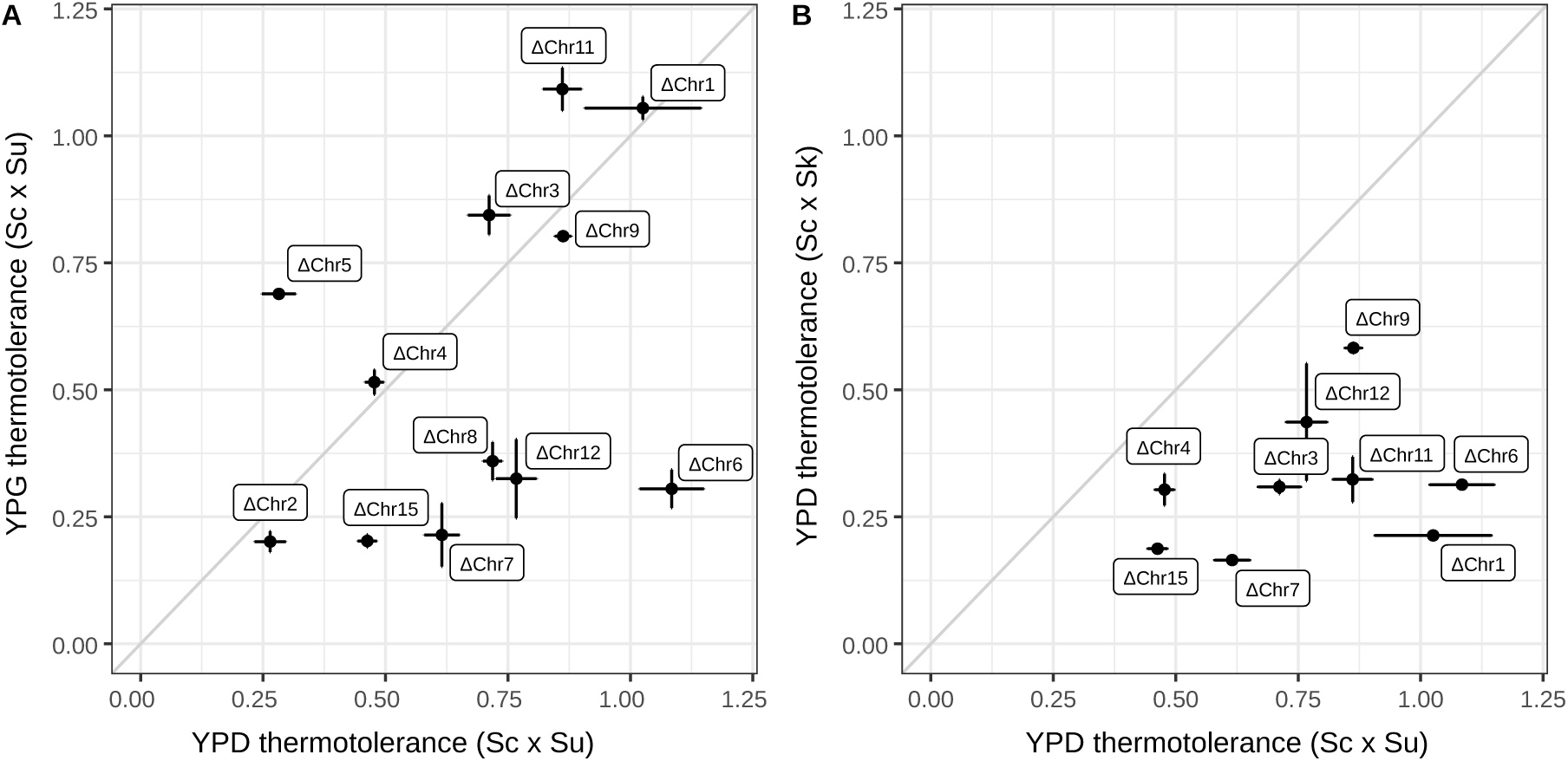
Thermotolerance of *S. cerevisiae* chromosome loss on different substrates and in different hybrid backgrounds. Thermotolerance was measured as the ratio of normalized mean colony size at 37 °C to 30 °C after three days of growth. Error bars represent standard error of the mean. (A) Thermotolerance of chromosome loss in *S. cerevisiae* x *S. uvarum* hybrids on dextrose (YPD) versus glycerol (YPG). (B) Thermotolerance of chromosome loss in *S. cerevisiae* x *S. uvarum* hybrids versus *S. cerevisiae* x *S. kudriavzevii* hybrids on dextrose (YPD). ΔChr 3 and ΔChr 12 *S. cerevisiae* x *S. kudriavzevii* strains are petite and ΔChr6 *S. cerevisiae* x *S. kudriavzevii* strains lost Chr 3R from *S. kudriavzevii*.

Given the large effects of *S. cerevisiae* chromosome loss in hybrids with *S. uvarum*, we also examined their effects in an *S. cerevisiae* x *S. kudriavzevii* hybrid. As *S. kudriavzevii* is less thermotolerant than *S. uvarum* [25], the thermotolerance effects of chromosomes could differ from those measured in the *S. cerevisiae* x *S. uvarum* hybrid. We generated a panel of *S. cerevisiae* x *S. kudriavzevii* hybrid chromosome loss strains for nine out of the sixteen *S. cerevisiae* chromosomes. All chromosomes except for Chr 12 showed temperature dependent growth defects (Table S2) and these effects were larger in the *S. cerevisiae* x *S. kudriavzevii* background compared to that of *S. cerevisiae* x *S. uvarum* (Figure 3B). However, this panel of strains had some heterogeneity in hybrid parental genotypes (Table S1) as well as phenotypes (Figure S1C, Figure S1E). Chr 3 and Chr 12 marked parental hybrids and their chromosome loss derivatives were petite and those with marked Chr 6 had lost the right arm of *S. kudriavzevii* Chr 3. Loss of the remaining six *S. cerevisiae* chromosomes showed thermotolerance effects that depend on the hybrid species context (Figure 3B).

Multiple *S. cerevisiae* chromosomes made large contributions to the thermotolerance of both interspecies hybrids. To quantify large thermotolerance effects we used a cutoff of 50% or more reduction in normalized growth at high compared to low temperature. On dextrose, there were four large effect chromosomes in hybrids with *S. uvarum* (2, 4, 5 and 15, Table S2) and seven in hybrids with *S. kudriavzevii* (1, 3, 4, 6, 7, 11, and 15, Table S2). On glycerol, there were six large effect chromosomes in hybrids with *S. uvarum* (2, 6, 7, 8, 12 and 15) and six in hybrid with *S. kudriavzevii* (1, 4, 6, 7, 11 and 15). This illustrates that large chromosomal effects are present across substrates and the hybrid background they are measured in.

### Multiple *S. cerevisiae* genes on chromosome 5 are required for thermotolerance

Loss of thermotolerance in the *S. cerevisiae* x *S. uvarum* chromosome loss strains could be due to recessive *S. uvarum* genes that are unable to properly function at high temperatures. However, the euploid hybrid does not grow as well as *S. cerevisiae* at high temperatures [29] indicating that *S. uvarum* genes may not be completely recessive. To examine dominance more carefully, we chose Chr 5, as it is collinear between species and shows a large thermotolerance effect on dextrose. To avoid any confounding influence of the *URA3* marker, we generated strains with reciprocal loss of Chr 5 and compared these with their euploid parents which also lacked the *URA3* marker.

At 30 °C, loss of *S. cerevisiae* Chr 5 had no effect on dextrose and increased growth on glycerol in comparison to the euploid hybrid, whereas loss of *S. uvarum* Chr 5 caused slightly lower growth on both dextrose and glycerol (Figure 4, p < 0.05, Table S4). This demonstrates that the *S. uvarum* chromosome is capable of supporting growth at levels similar to the euploid hybrid. However, at 37 °C loss of *S. cerevisiae* Chr 5 caused a substantial reduction in growth on both dextrose and glycerol compared to both the euploid hybrid and loss of *S. uvarum* Chr 5 (p < 0.05, Table S4). In comparison to the euploid hybrid, loss of *S. uvarum* Chr 5 caused higher growth at 37 °C on dextrose but not glycerol, suggesting that *S. uvarum* temperature sensitive alleles are not entirely recessive. We also found that at 4 °C loss of *S. cerevisiae* Chr 5 improved growth whereas loss of *S. uvarum* Chr 5 reduced growth in comparison to the euploid hybrid (Figure S2). The results of the reciprocal chromosome hemizygosity analysis confirm the large temperature effects of Chr 5 and indicate that *S. uvarum* has temperature sensitive alleles that are not entirely recessive at high temperature.

**Figure 4:**
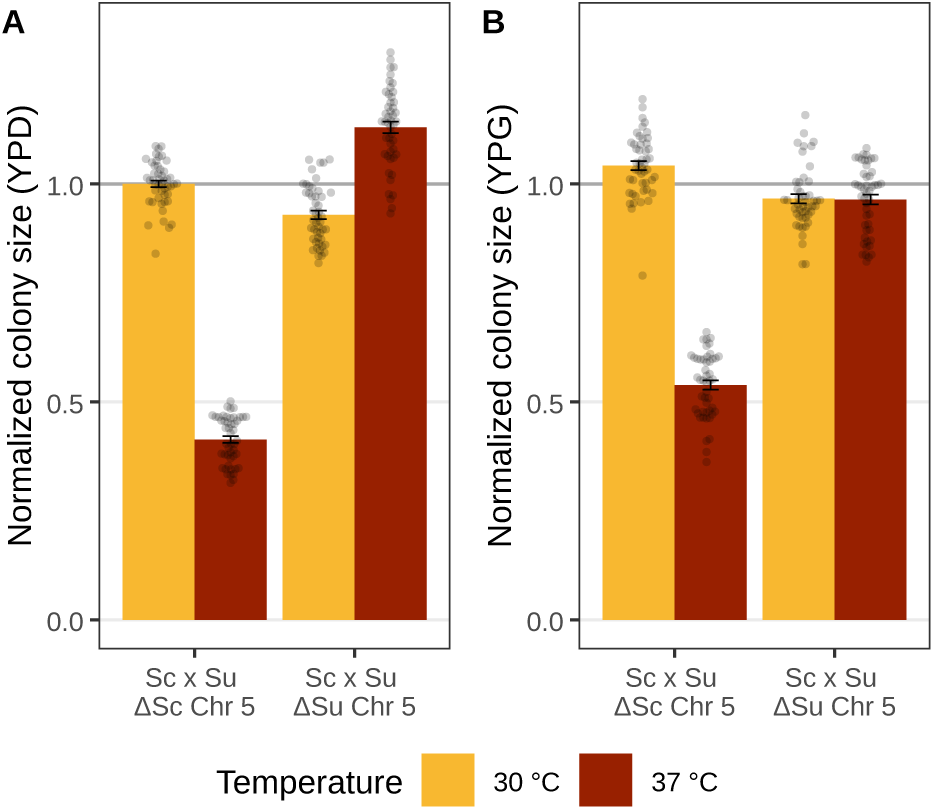
Thermotolerance of reciprocal chromosome 5 loss in *S. cerevisiae* x *S. uvarum* hybrids. Colony size after three days growth on dextrose (YPD) (A) and glycerol (YPG) (B) at 30 °C and 37°C. Both reciprocal hemizygous strains (Sc x Su ΔSc Chr 5 and Sc x Su ΔSu Chr 5) were normalized to colony size of a genotype matched euploid hybrid on respective substrates and temperatures. Error bars represent standard error of the mean and grey points are replicates.

To map thermotolerance on *S. cerevisiae* Chr 5 we generated a loss of heterozygosity (LOH) panel in the hybrid using CRISPR-Cas9 directed recombination [22]. CRISPR-induced double strand breaks (DSBs) along Chr 5 can result in partial or complete chromosome loss, or be repaired through mitotic recombination resulting in either gene conversion or break induced replication (Figure S3). Co-selection for loss of a telomeric *URA3* marker and retention of a centromeric *KanMX* marker enriched for break induced replication that generates LOH tracts extending from the DSB to the telomere or for partial loss of the target chromosome arm. Using five right arm and six left arm breaks, we isolated 82 LOH strains and sequenced their genomes (Table S5). Analysis of breakpoint distributions revealed clustering near the targeted CRISPR-Cas9 cut sites. Similar to a prior study [32], we also observed some breakpoints distant from the cut site (Figure S4), and for DSBs near the centromere we found breakpoints were distributed across the entire chromosome arm. One strain exhibited simultaneous partial LOH on both chromosome arms and three strains showed non-target aneuploidies (Table S5).

We quantified thermotolerance of the LOH panel and found both Chr 5 arms are required for thermotolerance. A centromere proximal 36 kb region of the left arm (5L) is required for high temperature growth on both dextrose and glycerol (Figure 5, Figure S5). On the right arm (5R) thermotolerance on dextrose but not glycerol was lost over a 203 kb region (302kb - 505kb) (Figure 5, Figure S5). Variance in thermotolerance was observed between some strains with similar breakpoints, particularly among strains with breakpoints close to Ty elements (Figure 5). One outlier strain from the 5L panel had LOH on both arms with both breakpoints near Ty elements (strain marked with * in Figure 5). These mapping results suggest the presence of at least one gene with a large to moderate effect on thermotolerance on 5L and multiple genes on 5R.

**Figure 5:**
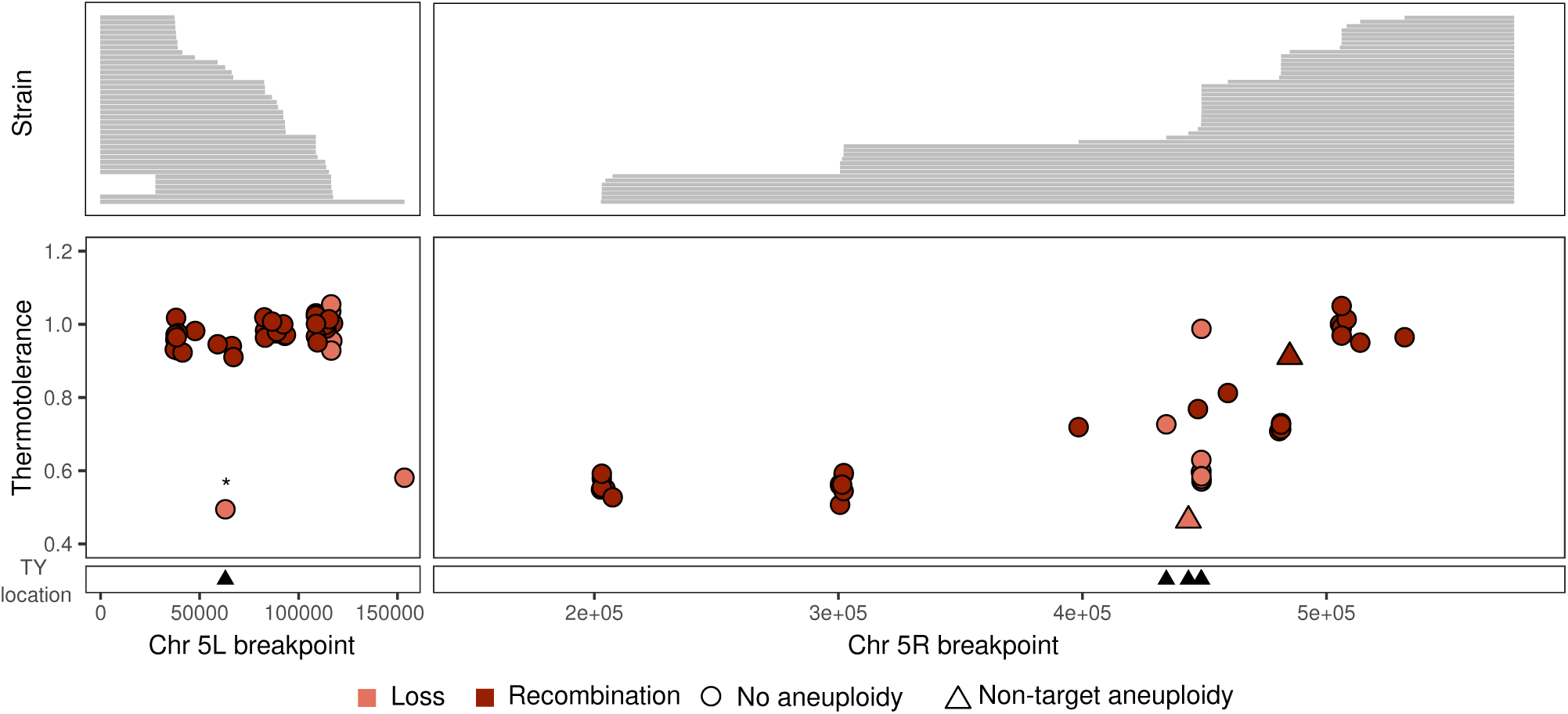
Thermotolerance of the Chr 5 loss of heterozygosity panel. Thermotolerance was measured by the ratio of normalized colony size at 37 °C to 30 °C on dextrose after three days of growth. Panels are split by the left arm (left) and right arm (right) of Chr 5. Upper panels indicate parts of *S. cerevisiae* Chr 5 that are absent (grey) corresponding to the breakpoints indicated on the x axis. In the lower panel, strains with non-target aneuploidy are represented by triangles, strains that showed recombination with *S. uvarum* Chr 5 are red, and strains with loss of *S. cerevisiae* Chr 5 segments without *S. uvarum* recombination are salmon. The strain marked with * lost parts of both 5L and 5R. Strains with breakpoints near Ty elements are marked on the x axis with black triangles.

### Chromosome 5L thermotolerance maps to *MMS21*

To identify *S. cerevisiae* genes responsible for Chr 5 thermotolerance we used the MoBY ORF plasmids carrying *S. cerevisiae* genes for complementation [33]. We tested 16 genes from the 36 kb interval on 5L and 21 genes on 5R from a ∼70 kb region (434kb to 505kb) where the drop in thermotolerance appeared linear (Table S6). Each plasmid was transformed into a strain with LOH spanning the region of interest on 5L or 5R and into the hybrid parent as a control for plasmid burden [34]. To minimize confounding effects of growth temperature on effect sizes, we measured colony sizes across multiple temperatures and calculated the temperature at which growth was inhibited to 50% of the maximum colony size (IT50). Out of the 16 genes we screened on 5L, only *MMS21* restored thermotolerance to control levels (Figure S6, Table S6). For the 21 genes screened on 5R, four candidates restored thermotolerance to control levels (IT50 difference < 1 °C), but these controls had lower thermotolerance relative to other transformant controls (Figure S6). Thus, we could not rule out temperature dependent effects caused by the plasmid alone.

We measured allele-specific effects of *MMS21* on thermotolerance using a reciprocal hemizygosity test. Deleting the *S. cerevisiae* allele of *MMS21* reduced thermotolerance (IT50) by 1.75 °C but there was no change in thermotolerance when the *S. uvarum* allele was deleted (Figure 6, Figure S7, t-test p < 2.2e-16). In this assay, the *MMS21* alleles account for 20% of the difference in thermotolerance between *S. cerevisiae* and *S. uvarum* (1.75/8.73). *MMS21* encodes a SUMO E3 ligase subunit of Smc5-Smc6 complex and was identified through a genetic screen for sensitivity to methyl methanesulfonate (MMS) [35]. We therefore tested *MMS21* alleles for effects on temperature and MMS sensitivity. We found that the *MMS21* reciprocal hemizygotes differed in their sensitivity to MMS at 33 °C or higher, but not at lower temperatures (Figure S8). Thus, the *S. uvarum* allele of *MMS21* acts like a temperature sensitive loss of function allele.

**Figure 6:**
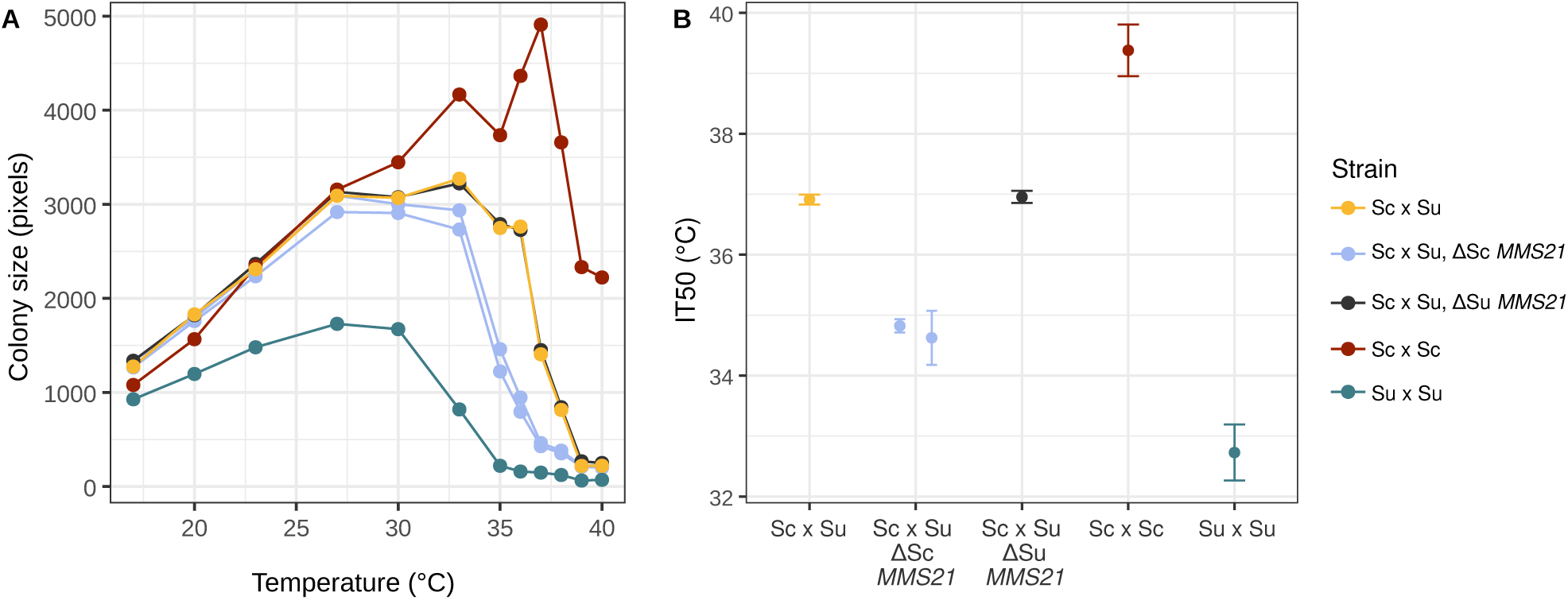
Thermotolerance of *MMS21* reciprocal hemizygotes. Thermotolerance was measured for *MMS21* hemizygotes in comparison to the hybrid parent, and *S. cerevisiae* and *S. uvarum* diploids on CM dextrose. Three transformation replicates were generated for *MMS21* hemizygotes, but only two ΔSc *MMS21* and one ΔSu *MMS21* strains (shown) had no other aneuploidy. (A) Mean colony size in pixels after two days of growth at various temperatures. (B) Temperature at which growth was inhibited by 50% (IT50). Points and error bars show strain means and standard deviation from a logistic regression.

### Deletion mapping of thermotolerance on Chr 5R

As plasmid complementation of genes on 5R did not conclusively yield any candidates, we generated deletions of multiple *S. cerevisiae* genes (ranging from 3 to 33 genes) across a ∼98 kb region (389kb to 487kb, Table S7) in the hybrid. Out of 46 deletions without any incidental aneuploidy, one strain (YJF5938) with a deletion of 17 genes (39kb) on *S. cerevisiae* 5R showed a substantial reduction in thermotolerance (Figure S9). Other deletion strains eliminated 16/17 genes absent in YJF5938, but none showed a similar reduction in thermotolerance. To test whether the one gene, *BEM2*, that was absent in YJF5938 but not the other deletion strains could account for the phenotype differences between the deletion strains and contribute to the effects of Chr 5 on hybrid thermotolerance, we generated *BEM2* reciprocal hemizygotes. Contrary to our expectation, deletion of *S. uvarum BEM2* resulted in a reduction in IT50 by 0.9 °C and deletion of *S. cerevisiae BEM2* did not significantly reduce IT50 (Figure S10). However, interpretation of these *BEM2* allele differences is confounded by chromosome aneuploidies present in the strains carrying the *S. cerevisiae BEM2* deletion.

## Discussion

Hybrid genetic analysis makes it possible to deconstruct trait differences between distantly related species that are reproductively isolated. We studied thermotolerance in hybrids of two *Saccharomyces* yeast species, *S. cerevisiae* and *S. uvarum,* that shared a common ancestor over 15 million years ago [36] and exhibit a rate of divergence at synonymous sites (1.08) [37] greater than that between human and mouse (0.73) [38]. Our study thus provides an important contrast to analysis of phenotypic divergence between closely related species. We find that hybrid thermotolerance depends on multiple *S. cerevisiae* chromosomes, and in the case of Chr 5 on both *MMS21* and gene(s) on the right arm. Below, we discuss our results in the context of how thermal divergence evolved and other factors relevant to interpreting the genetic basis of phenotypic differences between distantly related species.

### Multiple chromosomes of large effect

The genetic basis of phenotypic differences between species reflects the accumulation of genetic changes that occurred during divergence. Although recombination provides the ability to identify large to moderate effect loci it can be hard to reliably detect many small effects distributed across a chromosome [1,39]. Our chromosome loss panel does not resolve large and small effect loci within a chromosome, but it does have the advantage of being able to measure the combined impact of many genes in a single genetic background [40].

We find that most *S. cerevisiae* chromosomes have a large temperature effect in both *S. cerevisiae* x *S. uvarum* and *S. cerevisiae* x *S. kudriavzevii* hybrids. At face value this is inconsistent with the infinitesimal model, where the contribution of each chromosome is expected to be proportional to the number of genes on it: ranging from 1.8% (Chr 1) to 12.7% (Chr 4). While the large chromosomal effects could be caused by combined effects of many genes, we find two chromosomes (Chr 1 and Chr 11) with no detectable effect on either glucose or glycerol. However, the chromosomal effect sizes that we measure are based on the reduction in growth at high compared to permissive temperature and, as discussed below, this is likely an important consideration in interpreting small versus large effects.

Studies using chromosome substitution panels in mice also found multiple large effect chromosomes [41]. A central finding of these studies was that the sum of all chromosomal effects was much greater than the parental difference, implying strong epistatic effects. This result was found across multiple traits and for comparisons of both closely and more distantly related lineages [42]. This raises the possibility that our finding of multiple chromosomes of large effect may not be related to high divergence and may depend on how chromosomal effects were measured [43,44].

Phenotypes with a floor or ceiling can distort measurements of effect size [44]. In the context of thermotolerance, effect size may change as a function of temperature due to intrinsic temperature dependence. For example, a reduction in high temperature growth may depend on how much growth is present to begin with. We measured the effect of losing the *S. cerevisiae* allele of *MMS21* across temperatures and found that its effect was largest when the hybrid’s growth began to decline with temperature. The magnitude of its effect also changed with the difference in high temperature growth between the parent species (Figure 6). Empirical studies of fitness landscapes provide support for models of global epistasis, where a mutation’s effect size changes with the fitness of its genetic background [45,46]. In some instances similar relationships have been found between effect size and environmentally modulated growth rates [15,47]. These studies do not negate the importance of effect size on trait divergence but demonstrate how effect sizes can be distorted when viewed through the lens of phenotype assays designed to maximize the power of genetic analysis. In our study, both the genetic background and temperature at which the genetic changes historically occurred are unknown but relevant to whether or not evolution of thermotolerance occurred gradually over long periods.

### Multiple loci of large effect on chromosome 5

We found that both the left (5L) and the right (5R) arm of *S. cerevisiae* Chr 5 affect growth at 37°C on dextrose. These effects are not additive, as the phenotype resulting from complete loss of Chr 5 is only slightly more severe than the loss of either arm alone (Figure 4 versus Figure 5). *MMS21* appears to explain most of the Chr 5L effect because deletion of the *S. cerevisiae MMS21* allele caused colony size to be ∼40% of the euploid hybrid at 37 °C (Figure S7). For Chr 5R our data suggests the presence of multiple thermotolerance genes, and a small region of 5R near the centromere remains untested. However, there is some uncertainty in the mapping data as thermotolerance varied among strains with recombination breakpoints near Ty elements and we were unable to find any other aneuploidies that would explain this variability.

Our results do not exclude the possibility of many undetected genes on Chr 5 with small effects on thermotolerance. A previous study showed that replacement of multiple *S. cerevisiae* Chr 1 segments with homologous segments from *S. paradoxus* caused loss of thermotolerance [24]. In contrast, a genome-wide reciprocal hemizygosity screen in a *S. cerevisiae* x *S. paradoxus* hybrid only found one thermotolerance gene on Chr 1 [19,48]. Similarly, we found some disparity between chromosome segments and individual genes on Chr 5 in *S. cerevisiae* x *S. uvarum* hybrids. First, within the 70 kb interval associated with thermotolerance on Chr 5R we did not find any clear cases of complementation. However, there were seven genes in this interval that were missing in the MoBY plasmid library and were not tested. Second, deletion mapping provided evidence for an effect of *BEM2*, but we were unable to validate its effects using reciprocal hemizygosity analysis, in part due to associated aneuploidy. *BEM2* encodes a Rho GTPase activating protein that is required for bud emergence. While it is not essential, in *S. cerevisiae* deletion of *BEM2* results in defects in bud emergence and polarity, large and multinucleated cells, and loss of the ability to grow above 32 °C [49,50]. One explanation for the thermotolerance effects of Chr 5R is that they are caused by a combination of multiple *S. cerevisiae* genes with small or no effects on their own.

### *MMS21* divergence and thermotolerance

Reciprocal hemizygosity analysis showed that the *S. uvarum* allele of *MMS21* partially loses function at high but not low temperatures. This is unlikely to be caused by changes in gene expression as the *S. uvarum* allele of *MMS21* is expressed at higher levels than the *S. cerevisiae* allele in hybrids [51]. There are 50 amino acid differences between the two alleles that could influence protein function. *MMS21* encodes a SUMO E3 ligase, an essential component of the structural maintenance of chromosomes (Smc5-Smc6) protein complex that plays a crucial role in DNA repair and maintaining genomic stability. Loss of protein function could be mediated by temperature dependent disruption of *Mms21’s* interactions with the Smc5-Smc6 complex, or loss of SUMO domain function, as deletion of this domain results in reduced growth at high temperature [52]. SUMOylation targets of *Mms21* could have diverged between *S. cerevisiae* and *S. uvarum* resulting in altered post-translational profiles. It is also possible that divergence of *MMS21* is not directly linked to thermotolerance but related to conflict with selfish genetic elements. *MMS21* variants in *S. cerevisiae* have been linked to differences in 2µ plasmid stability, which is known to affect fitness [53]. While some *S. uvarum* strains carry extrachromosomal copies of 2µ plasmids, copies have also integrated into the nuclear genome [54]. Thus, *MMS21* divergence could affect high temperature growth via multiple routes.

### Evolution of thermotolerance in the *Saccharomyces* species

Distinguishing between the gain and loss of thermotolerance is important to inferring how thermotolerance evolved. A common observation for trait loss is that it can be independently caused by multiple genes either in the same lineage or across parallel lineages. Examples of such traits include freeze tolerance [55] and galactose metabolism [56] in the *Saccharomyces*. Under a trait loss model, once *S. uvarum* and *S. kudriavzevii* lost the ability to grow at high temperature, large effect temperature sensitive alleles would be free to accumulate across the genome. In contrast, trait gain often requires multiple coordinated changes. One way in which multiple genes of large effect could evolve in concert is if the effect size of each gene evolved gradually through several small effect changes. Another possibility is that genes with a large effect at high temperature cause smaller effects at lower temperatures, enabling gradual increases in thermotolerance over time. In either scenario, hybrid thermotolerance is expected to depend on multiple *S. cerevisiae* genes such that loss of one gene can cause loss of thermotolerance.

Our results show that in addition to the *S. cerevisiae* mitochondrial genome [20], multiple *S. cerevisiae* chromosomes are important for hybrid growth at high temperature. If gain of thermotolerance occurred along the *S. cerevisiae* lineage, we expect similar chromosomal effects on thermotolerance in *S. cerevisiae x S. uvarum* and *S. cerevisiae x S. kudriavzevii* hybrids. In contrast, if thermotolerance was lost independently in the *S. uvarum* and *S. kudriavzevii* lineages, we would expect hybrid specific patterns of chromosomal effects on thermotolerance. Our results indicate that while certain chromosomes consistently harbor thermotolerance loci across both hybrid backgrounds, the magnitude of their effects differs. *PAB1* (chromosome 5) and *DED1* (chromosome 15) have been previously implicated in thermal divergence between *S. cerevisiae* and *S. kudriavzevii* [57,58]. The *S. uvarum* allele of *PAB1* does not cause any detectable growth defects at high temperatures in our mapping data, suggesting that only *S. kudriavzevii PAB1* has lost the ability to support high temperature growth. Conversely, loss of *S. cerevisiae* chromosome 15 which harbors *DED1*, shows a large effect on thermotolerance in both hybrid backgrounds, consistent with a gain in thermotolerance along the *S. cerevisiae* lineage. Taken together, these observations support a model in which the *S. cerevisiae* lineage acquired thermotolerance while *S. uvarum* and *S. kudriavzevii* experienced independent, parallel loss in thermotolerance [26]. However, distinguishing between gain or loss of thermotolerance becomes less important if there were trade-offs in changes to the thermal performance curve [59,60].

Evolutionary tradeoffs between high and low temperature growth in *Saccharomyces* can place constraints on how temperature tolerance evolves. Both *S. uvarum* and *S. kudriavzevii* grow better than *S. cerevisiae* at low temperatures [25,29,61]. In *S. cerevisiae x S. uvarum* hybrids mitochondria type and specifically *COX1* shows a trade-off between growth at high and low temperature [20]. Evolution of hybrids at different temperatures resulted in loss of heterozygosity which favored temperature dependent retention of parental alleles [62].

Tradeoffs also exist in parental contexts, as seen with allele replacements of thermotolerance genes in *S. cerevisiae* with their *S. paradoxus* orthologs [63], and larger tradeoffs in experimental evolution of cold tolerant compared to heat tolerant Saccharomyces species [60]. In our study, we observe a tradeoff between high and cold temperature growth of the reciprocal chromosome 5 loss strains (Figure S2). The increasing evidence for temperature related tradeoffs raises the possibility that the loss of thermotolerance observed in *S. kudriavzevii* and *S. uvarum* could be a consequence of gain of cold tolerance in these lineages. The mechanism of cold tolerance in these species involves different transcriptional and metabolic networks [26,64,65] and could cause independent parallel loss of thermotolerance.

### The unique hybrid environment

One limitation of using hybrids for genetic analysis is that they represent a unique genomic and cellular environment. Due to differences in gene expression patterns and protein stabilities [27,29,51], the phenotypic effects of a change in hybrids may not be fully recapitulated in parental contexts. *S. cerevisiae* x *S. uvarum* hybrids can tolerate both high and low temperatures and are phenotypically distinct from the parental species [29]. The thermotolerance effect of chromosomes could vary between hybrids and parental backgrounds. For example, substitution of *S. uvarum* chromosome 5 into a *S. cerevisiae* background is not viable unless chromosome 7 is replaced as well, suggesting that epistatic interactions may influence viability [66]. While there are examples of alleles affecting thermotolerance in both hybrid and parental contexts, their effect size can differ [19]. Reciprocal allele replacements of thermotolerance genes between the two species will be valuable to understanding the differences between hybrid and parental backgrounds and interpreting hybrid mapping studies.

### Conclusions

Our study provides insight into the genetic basis of phenotypic divergence over long evolutionary timescales by taking advantage of hybrid genetic analysis. Few studies have mapped genes underlying trait differences between species that diverged more than 5 million years ago [11]. Notable examples include the identification of loci underlying floral scent and coloration differences between Petunia species, as well as Mimulus species [67–70]. Our results show how hybrids can make highly diverged species accessible to genetic analysis and that thermotolerance in the *Saccharomyces* depends on multiple loci. However, further work is needed to determine how these loci affect thermotolerance in their parental context and whether they evolved through single or multiple substitutions.

## Methods

### Chromosomal mapping

#### Strains and growth conditions

Sixteen haploid *S. cerevisiae* strains with *URA3* and *KanMX* markers flanking the centromere of each chromosome [30](Table S7) were crossed to *S. uvarum* (YJF2562) or *S. kudriavzevii* (YJF2801) to generate hybrids. *MAT* PCR and mitotype PCR were used to select hybrid colonies with *S. cerevisiae* mitotypes. All strains were grown at 25 °C in YPD (1% yeast extract, 2% peptone, and 2% dextrose) unless otherwise specified.

#### Generating hybrid strains missing the target S. cerevisiae chromosome

The sixteen centromere marked *S. cerevisiae* x *S. uvarum* hybrids (YJF3620 - YJF3635) were plated on CM with 1 mg/mL 5-fluoroorotic acid (5-FOA)(CM: 0.3% yeast nitrogen base with amino acids, 0.5% ammonium sulfate, and 2% dextrose) to select against the *URA3* marker on the target *S. cerevisiae* chromosome. Once colonies were observed (3-5 days of growth), they were re-streaked on 5-FOA plates and single colonies were isolated and patch tested on YPD supplemented with G418 (200 mg/L) to test whether the *KanMX* marker was still present. Three to five colonies of strains that lost the *KanMX* and the *URA3* marker were recovered for twelve out of the sixteen hybrids. Hybrid chromosome loss strains missing *S. cerevisiae* chromosome 10, 13, 14 or 16 could not be generated. The same procedure as described above was used to generate *S. cerevisiae* x *S. kudriavzevii* chromosome loss strains. Strains missing chromosome 2, 5, 8, 10, 13, 14, 15, or 16 could not be generated.

#### Whole genome sequencing and analysis

To confirm loss of the target *S. cerevisiae* chromosome and to account for any other non-target aneuploidy, we performed whole genome sequencing on the twelve *S. cerevisiae* x *S. uvarum* and nine *S. cerevisiae* x *S. kudriavzevii* hybrid parental strains, and all the strains derived from them (Table S7). DNA was extracted using the Quick-DNA Fungal/Bacterial Microprep kit from Zymo. The Nextera DNA Library Preparation kit (Illumina) was used to make paired end libraries and genomes were sequenced on a Illumina Hiseq X (2 x 150 bp) by Novogene.

Reads were mapped to either a *S. cerevisiae* x *S. uvarum* hybrid reference genome created by concatenating *S. cerevisiae* (S288C-R64-2-1) and *S. uvarum* reference genomes [71], or to a *S. cerevisiae* x *S. kudriavzevii* hybrid reference genome created by concatenating *S. cerevisiae* (R64-4-1) and *S. kudriavzevii* (CR85, GCA_003327635.1) reference genomes using bowtie-2 [72]. Duplicates were marked and removed with picard tools 2.12.0 [73]. Coverage at each position was calculated using samtools depth [74]. We calculated a rolling average depth over 1,000 bp windows with 500 bp increments for each chromosome. These plots were manually inspected and genotype calls were recorded in Table S1.

#### Phenotyping

*S. cerevisiae* x *S. uvarum* hybrids*, S. cerevisiae* x *S. kudriavzevii* hybrids, and their the derivatives were arrayed in a 1,536 format in a randomized order on rectangular 2% YPD agar plates (number of technical replicates ranged from 8 to 12). After two days of growth, this master plate was replica plated onto phenotyping plates using the Singer Rotor HDA (Singer Instruments, UK). Phenotyping was done on two substrates, YPD (fermentable) and YPG (respiratory, 1% yeast extract, 2% peptone, and 3% glycerol) at 30 °C and 37 °C (three replicate plates for each temperature). Plates were incubated at the respective temperatures and imaged for four days using the Phenobooth (Singer Instruments, UK) at a resolution of 4,128 × 3,096. Colony size was measured in pixels using the Phenosuite software v.2.22.0607.1. Colony sizes from day three images were used for further analysis.

Spotted positions lacking detected colony size despite visible growth, likely due to overgrowth of colonies, were excluded from the dataset. We evaluated the use of row-column normalization to control for potential edge effects, following the method described by Miller et al. [75]. However, this normalization procedure did not improve the r² for a linear model fitting colony size to genotype. Consequently, no normalization for edge effects was applied to the final dataset. To compare across different growth conditions (temperature, substrate) and strains, colony sizes were normalized by dividing the mean colony size of the chromosome loss strain by the mean colony size of the corresponding hybrid parent strain grown under identical substrate and temperature conditions. To assess whether the loss of a target chromosome resulted in a temperature-specific growth phenotype, paired t-tests with Bonferroni correction were performed comparing the normalized colony sizes at 37 °C to those at 30 °C for each chromosome loss strain.

### Chromosome 5 loss of heterozygosity panel

#### Reciprocal chromosomal loss of heterozygosity for chromosome 5 in S. cerevisiae x S. uvarum hybrids

To generate the reciprocal loss and the loss of heterozygosity panel along *S. cerevisiae* chromosome 5, we first generated strains with *URA3* on chromosome 5. For construction of these strains, transformations were carried out according to the standard lithium acetate protocol [76] with a minor modification: after a 30 minute incubation at room temperature, the transformation reactions were heat shocked for 1 hour at 37 °C. Transformants were confirmed by junction PCR. To avoid deficiencies in laboratory strains we used a derivative of an oak tree strain YPS163 as the starting strain for *S. cerevisiae* and a derivative of CBS7001 for *S. uvarum*. First, the endogenous *URA3* gene in both these strains was replaced with a *KanMX* cassette flanked by LoxP sites. These strains were then transformed with a plasmid carrying Cre recombinase (pSH47) to eliminate the *KanMX* cassette (YJF4559 and YJF4581). Next, *URA3* was inserted at the telomere of *S. cerevisiae* chromosome 5L in the intergenic region between *HPA3* and *SIT1* (YJF4583). We also inserted a *KanMX* marker near the centromere, upstream of *MNN1* on *S. cerevisiae* 5R (YJF5098). This *S. cerevisiae* strain was crossed to a *rho^-^* version of *S. uvarum Δura3* (YJF4682) to generate an inter-species hybrid strain with *URA3 on S. cerevisiae* chromosome 5L (YJF5192). Similarly, a *URA3* marker was inserted at the telomere of chromosome 5R between *PUG1* and YER186C and a *KanMX* marker was inserted near the centromere, upstream of *MNN1* in the *S. cerevisiae* strain YJF4559 to obtain YJF5207. Crossing this strain to *S. uvarum* generated a hybrid with *URA3* at the telomere of *S. cerevisiae* chromosome 5R (YJF5216).

To generate reciprocal loss of *S. uvarum* chromosome 5 in a hybrid background, a *URA3* marker was inserted at the telomere of chromosome 5L in the *S. uvarum Δura3* strain (YJF4581) in the intergenic region between *HPA3* and *SIT1* (YJF5173). This strain was then crossed to *S. cerevisiae* strain YJF4559 to generate a hybrid with *URA3* on *S. uvarum* chromosome 5 (YJF5203). The two hybrid strains YJF5192 and YJF5203 were grown overnight in YPD at 25 °C and plated on CM with 5-FOA (1 mg/mL) to select for loss of the marked chromosome 5. This resulted in two hybrid strains, one missing *S. cerevisiae* chromosome 5 (YJF5211) and another missing *S. uvarum* chromosome 5 (YJF5212). Whole genome sequencing of these two strains confirmed loss of the target chromosome and they both carry *S. cerevisiae* mitochondria.

#### Generating a loss of heterozygosity panel along S. cerevisiae chromosome 5 in the hybrid background

To generate varying tracts of LOH along *S. cerevisiae* chromosome 5 in the hybrid background, a method previously used by Sadhu et. al [22] was modified as described below. CRISPR Cas9 plasmid pML104-HygMx4 (Addgene plasmid # 83475 from Patricia Wittkopp) was used to target five double strand breaks (DSBs) along *S. cerevisiae* chromosome 5L and six DSBs along *S. cerevisiae* chromosome 5R. Guide RNA oligos were designed and cloned into pML104-HygMx4 as described in Laughery et. al [77]. Oligos and target sequences are listed in Table S8. All guide RNAs target intergenic regions and are specific to *S. cerevisiae* chromosome 5. Next, CRISPR plasmids that target chromosome 5L were transformed into hybrid strain YJF5192 and plasmids that target chromosome 5R were transformed into YJF5216. Transformation reactions were allowed to recover in liquid YPD for 4 hours and then grown overnight in YPD supplemented with hygromycin B (200 mg/L) to select for transformants with the CRISPR plasmid. The next day these cultures were spun down, and re-suspended in 200 μL of water and plated on CM with 5-FOA (1 mg/mL) and G418 (200 mg/L) to enrich for cells that repaired the Cas9 induced DSBs by break induced replication and select against aneuploids and gene conversion (Figure S3). Colonies were re-streaked to isolate single genotypes and eight colonies for each cut site were sequenced and phenotyped.

#### Whole genome sequencing and data analysis

To identify breakpoints and the extent of LOH along *S. cerevisiae* chromosome 5 in the hybrids, the LOH panel and the parental hybrid genomes were sequenced. DNA was extracted using the high throughput Bio-On-Magnetic-Beads [78] protocol with modifications. After strains were grown to saturation in YPD cultures, 200 μL was transferred to 96 well plates and centrifuged for 5 minutes at at 1,000 xg. After removing the supernatant, the cell pellets were resuspended in 100 μL lyticase buffer (1.2M sorbitol, 50 mM EDTA, 2.5 units Zymolase, 30 μg RNaseA) and incubated at 37 °C for 30 minutes. The resulting spheroplasts were then collected by centrifugation as above and resuspended in 60 μL of lysis buffer (4M guanidinium thiocyanate, 2% w/v N-lauroylsarcosine sodium salt, 20 mM EDTA, 50 mM Tris-HCl pH 8, adjusted to pH 7.6 with HCl) and incubated at room temperature for 5 minutes. 35 μL of bead buffer (20% w/v PEG 8000, 2.5 M NaCl, 25 mM EDTA pH 8.0) and 25 uL AmpureXP beads (Beckman Coulter Life Sciences) were added to the lysates and mixed by pipetting. After incubation on a magnetic stand for two minutes followed by removal of the supernatant, the beads were washed thrice with 80% v/v ethanol and any remaining ethanol was removed by pipetting. After a minute incubation at room temperature to allow the remaining ethanol wash to evaporate, the plate was removed from the magnetic stand and beads were resuspended in 25 μL of DNAse-free water pre-warmed to 56C. The plate was incubated on the magnetic stand for 5 minutes and the supernatant with the extracted DNA was transferred to a new plate. Libraries were prepared using the Nextera DNA Flex Library Preparation kit from Illumina and sequenced using the NovaSeq Paired-end 150 platform at Novogene. Reads were mapped to an *S. cerevisiae* x *S. uvarum* hybrid reference genome created by concatenating *S. cerevisiae* YPS128 assembly (GCA_002057995.1) and the *S. uvarum* YJF1449 assembly (PRJNA882904) using bowtie-2 [72]. Coverage analysis was conducted as described above. Recombination breakpoints were manually identified and annotated by analyzing the sam files in the Integrative Genomics Viewer [79].

#### Phenotyping

The LOH panel, the parental hybrids and genotype matched hybrid YJF5334 were randomized on a 1,536 rectangular YPD plate. Phenotyping was carried out as described above in the Chromosomal mapping section. Plates incubated at 4 °C were imaged at day 50. To control for edge effects, row - column normalization was carried out as described in Miller et. al [75]. In this instance, normalization improved the r^2^ for a linear fit between Genotype and colony size. Normalized colony size was calculated as described above using the hybrid YJF5334 as the control for the entire 5LOH panel.

### MoBY plasmid screen

Frozen bacteria stocks containing selected MoBY plasmids (Table S6)[33] were resurrected on LB agar plates containing 5 mg/L tetracycline, 12.5 mg/L chloramphenicol and 100 mg/L G418 incubated at 30 °C. Cultures started from single colonies were grown at 30 °C with 300 rpm shaking in 2 mL LB containing the same antibiotics and harvested 24 hours later for plasmid extraction by MiniPrep kit (Qiagen). Hybrid (*S. cerevisiae* x *S. uvarum*) strains GS6_A1 and YJF5340 were transformed using the protocol described in Gietz et al [76] with modifications. YPD cultures (50 mL) were harvested at mid-log (OD600 ∼ 1) by centrifugation at 1,000 xg for 2 minutes followed by 2 washes with equal volumes of sterile water. Cell pellets were resuspended in 10 mL 100 mM lithium acetate and incubated at room temperature for 20 minutes followed by centrifugation at 1,000 xg for 1 minute. Cells were resuspended in a 10.5 mL mixture of 100 mM lithium acetate, 40% w/v polyethylene glycol 3500, and 4% v/v salmon sperm DNA, and distributed into a 96-well plate in 100 μL aliquots. 10 μL plasmid DNA (∼50-500 ng) was added to each well and mixed by pipetting. After a 30 minute incubation at room temperature, the plate was incubated at 37 °C for 1 hour followed by a 30 second spin at 1,000 xg to pellet the cells. The supernatant was removed by pipetting and cells were resuspended in 300 μL YPD and incubated at 23 °C overnight. Cultures were plated on either YPD agar containing 200 mg/L G418 or CM-uracil agar (0.5% ammonium sulfate, 0.17% yeast nitrogen base, 0.2% dropout mix without uracil, Table S6) for viable colonies transformed with plasmids containing left or right arm genes, respectively. Plates were incubated for 3 days at 23 °C from which transformants were picked for thermotolerance profiling by solid agar growth assays.

#### Phenotyping

Thermotolerance was assessed by colony size measurements on agar plates incubated at various temperatures (23, 30, 34, 35.5, 37, 38, 39 °C). Individual colonies from each transformation (n ∼ 6-12) were arrayed on 2% agar plates with corresponding media and selection in a 384 colony layout using a previously described method [75]. Printed plates were grown at 23 °C for 2 days and replica plated to corresponding media and selection for 3 technical plating replicates at each test temperature using a Rotor HDA robot. Plate images were captured at 2,560 X 1,920 pixels every day for 5 days at the indicated incubation temperatures using a Phenobooth. The temperature at which 50% of the maximum colony size would be observed (IT50) was predicted using 5 point logistic regression through the nplr package (v0.1-8) in R [80]. In order to maximize the regression fit through the measured points in the slope, colony sizes for a given sample that were smaller at lower temperatures than higher temperatures were adjusted to match the highest measured size observed at any of the higher temperatures so that the resulting curves start with the maximum colony size observed followed by decreasing sizes at higher temperatures as observed.

### Deletion mapping on chromosome 5R in hybrids

#### Deletion of chromosome 5R regions

To map thermotolerance within *S. cerevisiae* 5R we generated a hybrid mapping panel with deletions of varying lengths along this arm. To generate the panel, primers were designed with 40bp overlaps to intergenic regions unique to *S. cerevisiae* Chr 5R (primers listed in Table S8) and 20bp overlap with the *URA3* cassette. Eleven sets of forward/reverse primers were designed to generate overlapping deletions of ∼10 kb segments across the 97 kb region. In addition, all possible combinations of the same set of primers (n = 74) were used to generate deletions of varying sizes (725 bp-97kbp) across the entire region. We transformed the euploid hybrid (YJF5550) with these templates and screened URA+ transformants by PCR first and then by whole genome sequencing (described in the loss of heterozygosity panel section above). Out of 360 strains, 55 were confirmed to carry a deletion on Chr 5R and 246 had no other incidental aneuploidy in the remainder of the genome. Together, we recovered 46 deletions without other aneuploidies, including one partial deletion that did not span the entire intended region (YJF6065, Table S7).

#### Phenotyping

To measure thermal phenotypes of the deletion mapping panel, colony sizes of genotype-confirmed strains were determined from plates incubated at various temperatures (23, 28, 30, 33, 35, 36, 37, 38, 39, 40 °C). First, the deletion mapping panel was arrayed on CM plates in 96 colony arrays using the methods described elsewhere [75]. Colonies were allowed to grow at 23 °C for 2 days and then replica plated to 3 technical replicate plates at each test temperature using a Rotor HDA robot. Next, daily images of the phenotyping plates were captured at 2,560 X 1,920 pixels for four consecutive days using a Phenobooth. For consistency with the Chr 5 LOH mapping, we measured thermotolerance by the ratio of growth at 37 to 30 °C after three days and then normalized to the euploid control.

### *MMS21* and *BEM2* reciprocal hemizygosity tests

Species-specific *MMS21* knockouts were constructed in the hybrid *S. cerevisiae* x *S. uvarum* strain YJF5550 using allele replacement with the *URA3* cassette via homologous recombination (Table S7). *URA3* was amplified from plasmid pML104 (pML104 from John Wyrick, Addgene plasmid # 67638) using primers with 40 bp overlap to flanking regions of *S. cerevisiae* or *S. uvarum MMS21* (Table S8). PCR products were cleaned using a QIAquick PCR Purification Kit (Qiagen, USA) and transformations into YJF5550 were carried out as described in the MoBY plasmid screen section using CM-uracil agar to select for transformants. Insertion of *URA3* into the intended locus was confirmed by PCR. Three confirmed knockouts of each allele were used for thermotolerance assessment by colony size assay. After whole genome sequencing, one strain with an *S. cerevisiae MMS21* deletion and two strains with an *S. uvarum MMS21* deletion were removed due to incidental aneuploidies elsewhere in the genome. For *BEM2*, knockouts were made in *S. cerevisiae* and *S. uvarum* parental strains separately, and then mated to parental controls to generate reciprocal hemizygotes (see Table S7 for deletion strains and Table S8 for primers).

#### Phenotyping

For phenotyping, strains were arrayed in a snaking pattern in a 96 colony configuration (n = 8 per strain) and printed on CM agar plates. After 2 passages on CM agar grown at 23 °C for 3 days each, the strains were printed on CM agar plates using a Singer Rotor HDA and incubated at various temperatures 20, 23, 27, 30, 33, 35, 36, 37, 38, 39 and 40 followed by colony size measurements using a Phenobooth and IT50 calculations using the methods described in the MoBY plasmid screen section. For *MMS21* strains, colony size response to temperature and methyl methanesulfonate (MMS) were assessed at 23, 30, 33, 35, 37 °C with 0 to 0.05% vol/vol MMS added to the molten CM agar solution.

## Acknowledgments

We thank Jun-Yi Leu for sharing the centromere marked *S. cerevisiae* strains, Paul Sniegowski for the *S. cerevisiae* oak strain, Amparo Querol for the *S. kudriavzevii* strain, Chris Hittinger for sharing the *S. uvarum* strain and Patricia Wittkopp and John Wyrick for sharing plasmids. We thank Tiffany Dias and Timothy Lai for laboratory assistance and members of the Fay lab for comments and feedback on the project. This work was supported by NIH grant GM080669 and GM156247.

## Data Availability

Raw phenotyping data and code for figures can be found at 10.60593/ur.d.34018224, accessible prior to publication at https://figshare.com/s/4ab1f2d168763d196e18. Sequencing data was deposited at NCBI under BioProject PRJNA1512722.

