## Supplementary material for "Deconstruction of *Saccharomyces* hybrids reveals multiple genomic regions required for thermotolerance": Figures S1-S10

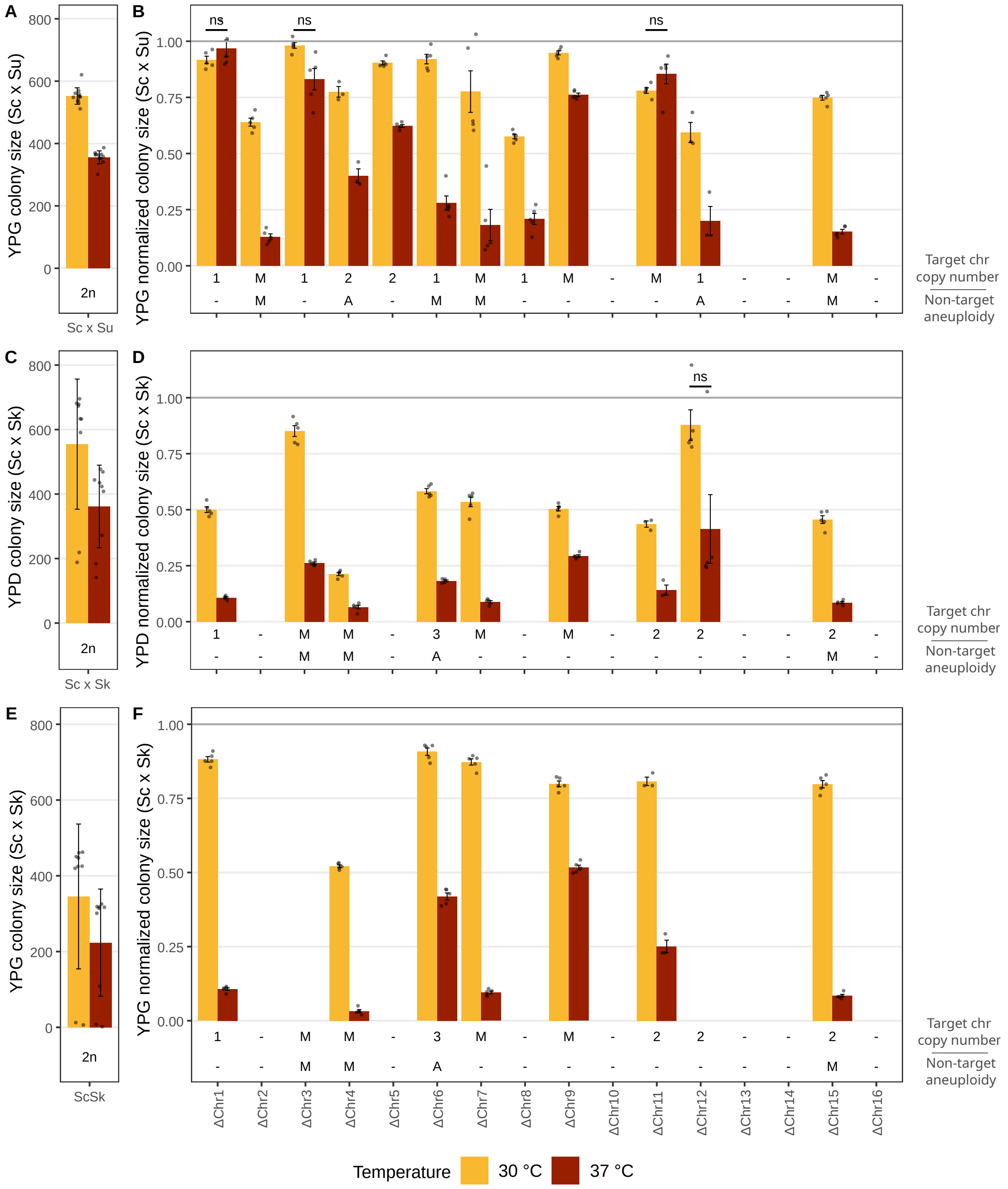


Figure S1: Thermotolerance of parental hybrids and chromosomal loss strains. Thermotolerance was measured by colony size after three days of growth at 37 °C (red) versus 30 °C (yellow). A and B are *S. cerevisiae x S. uvarum* on glycerol (YPG), C and D are *S. cerevisiae x S. kudriavzevii* on dextrose (YPD) and E and F are *S. cerevisiae x S. kudriavzevii* on glycerol (YPG). In A, C, and E, bars and whiskers represent the mean and standard deviation. In B, D and F, bars and whiskers represent the mean and standard error of the mean. Grey points are individual strains. All chromosomes except those marked with ns had significant thermotolerance effects ($p < 0.05$, Table S2). Annotations below the plot indicate the copy number of the target chromosome (First line: 1, 2, 3 or M for mixed) and the presence of non-target aneuploidy (Second line: A = present, M = mixed).

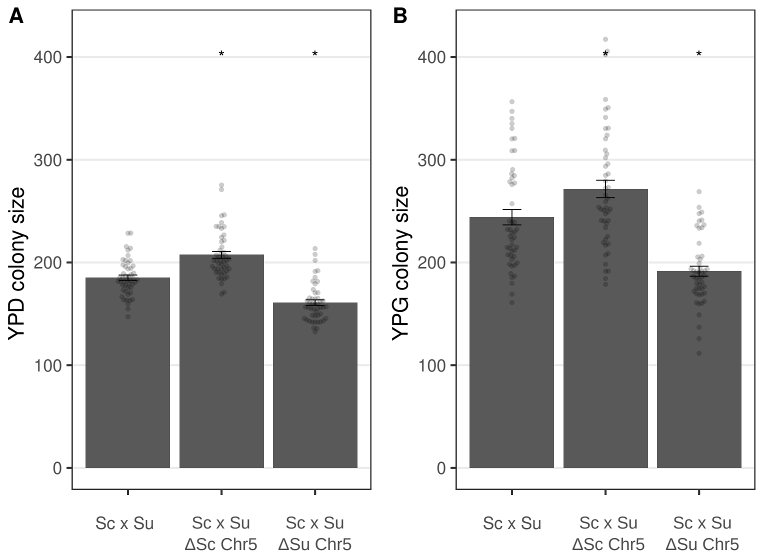


Figure S2: Cold tolerance of the euploid hybrid and chromosome 5 reciprocal hemizygotes. Colony size was measured after 50 days growth at 4 °C on dextrose (A) and glycerol (B). Bars represent mean colony sizes and bars are standard error of the mean. Hemizygotes that significantly differ from the euploid hybrid are marked with *.

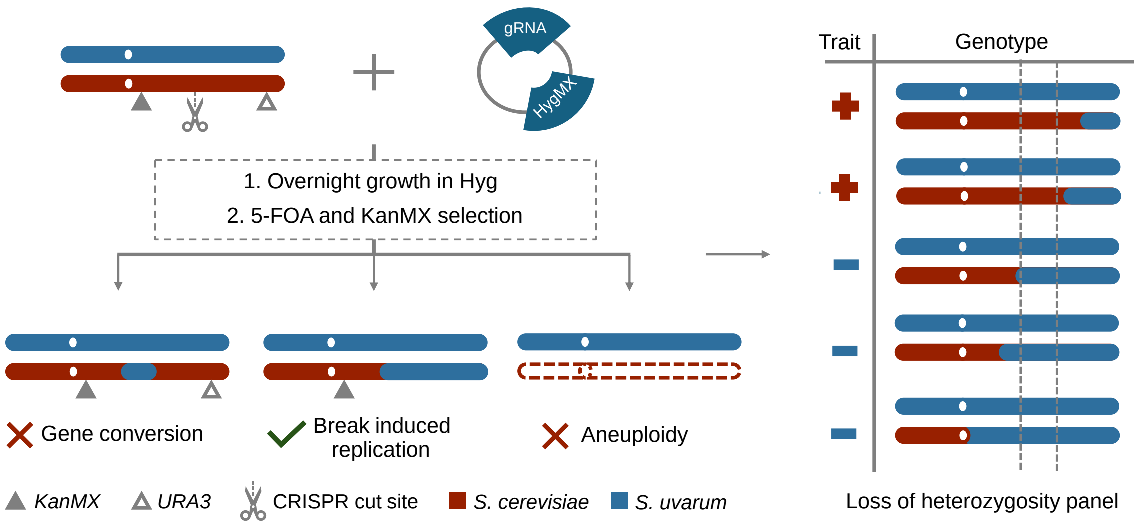


Figure S3: Loss of heterozygosity (LOH) outcomes and trait mapping. The hybrid strain with a centromeric *KanMX* marker and a telomeric *URA3* marker was transformed with Hyg CRISPR plasmids that cut the target *S. cerevisiae* chromosome and grown overnight with hygromycin selection. Transformation reactions were plated on 5-FOA and G418 double selection plates to simultaneously select for loss of the telomeric marker and presence of the centromeric marker. This excludes transformants where DSBs have been repaired by gene conversion or have completely lost the target chromosome and enriches for break induced replication events or partial chromosome loss without replication (not shown). Phenotyping of the resulting LOH panel maps a trait to genomic regions associated with LOH.

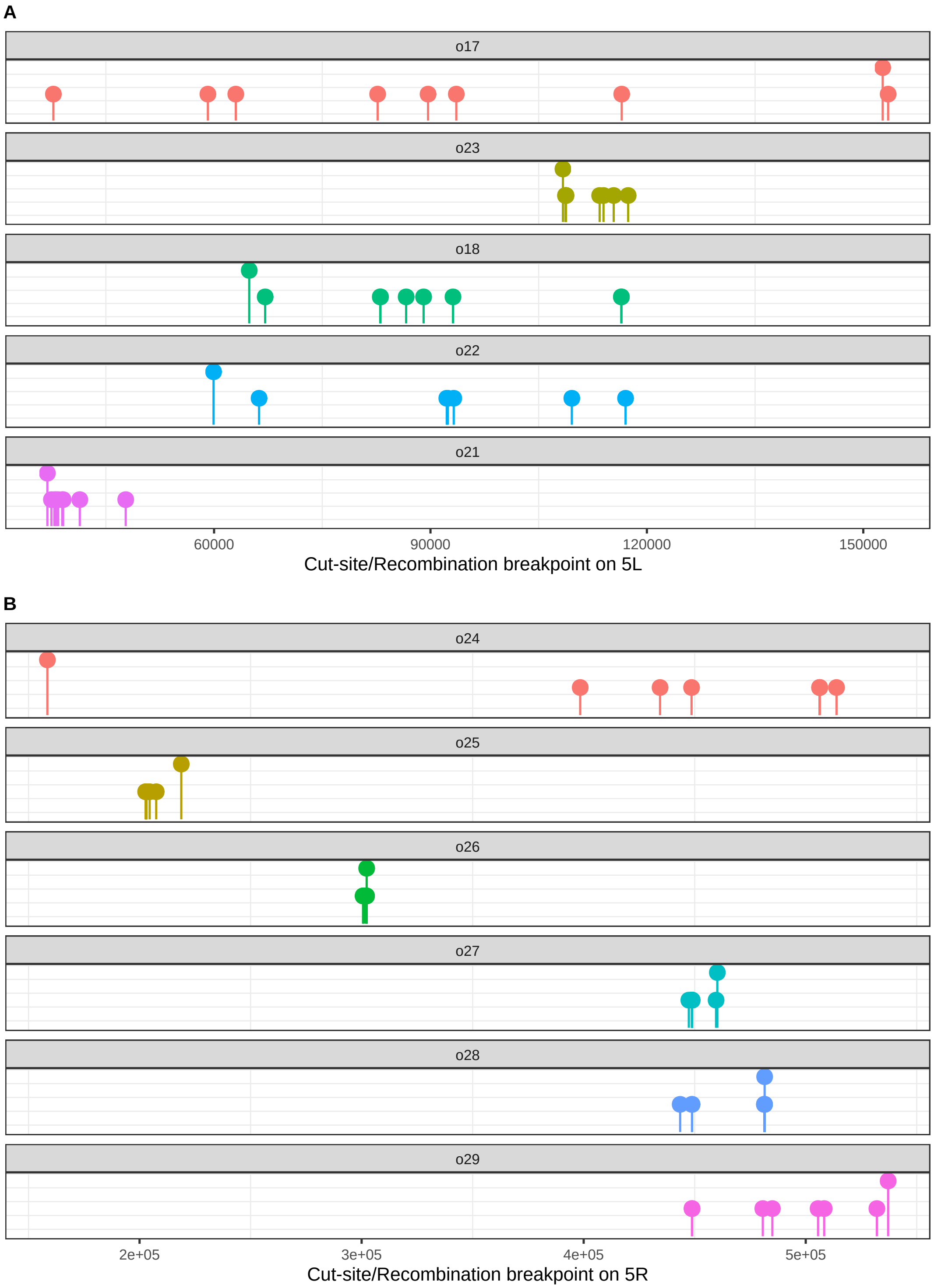


Figure S4: CRISPR cut sites and recombination breakpoint locations along *S. cerevisiae* chromosome 5. (A) Left arm. (B) Right arm. Raised points indicate CRISPR target sites for the respective guide RNA oligos (o17 - o29) and the shorter points are the recombination breakpoints in strains recovered post repair of the double strand breaks.

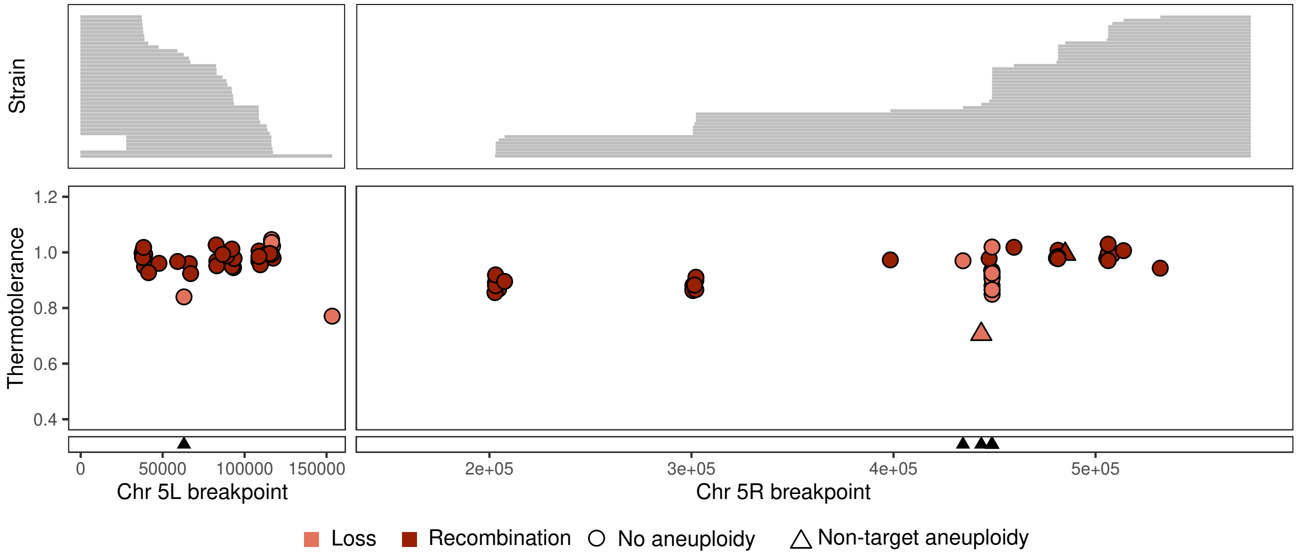


Figure S5: Thermotolerance of the Chr 5 loss of heterozygosity panel on glycerol. Thermotolerance was measured by the ratio of normalized colony size at 37 °C to 30 °C after three days of growth. Panels are split by the left arm (left) and right arm (right) of Chr 5. Upper panels indicate parts of *S. cerevisiae* Chr 5 that are absent (grey) corresponding to the breakpoints indicated on the x axis. In the lower panel, strains with non-target aneuploidy are represented by triangles, strains that recombined with *S. uvarum* Chr 5 by break induced replication are red, and strains with loss of *S. cerevisiae* Chr 5 segments without *S. uvarum* recombination are salmon. Strains with breakpoints near Ty elements are marked on the X axis with black triangles.

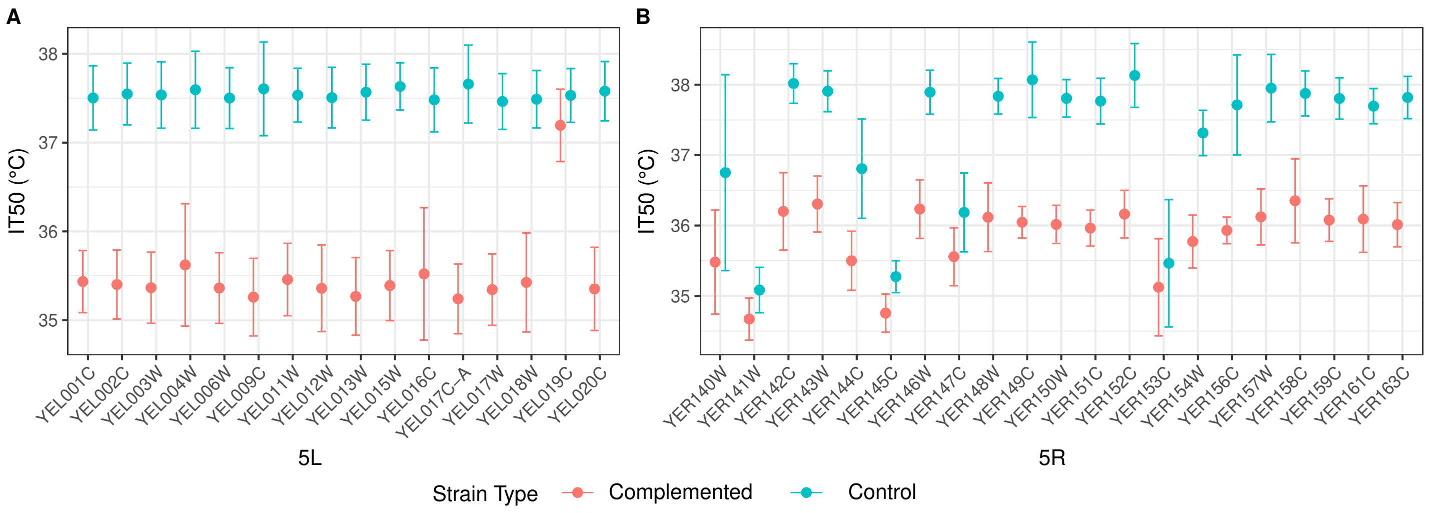


Figure S6: MoBY plasmid complementation screen. Thermotolerance after 2 days incubation of strains transformed with plasmids carrying *S. cerevisiae* genes from the left (A) or right (B) arm of Chr 5 as measured by the temperatures at which the maximal colony growth is inhibited by 50 percent (IT50) as predicted by 5 point logistic regression. Complemented strains lack either the left (GS6 A1) or right (YJF5340) arm of S. cerevisiae Chr 5 whereas control strains (YJF5334 and YJF5337) each have a fully intact Chr 5.

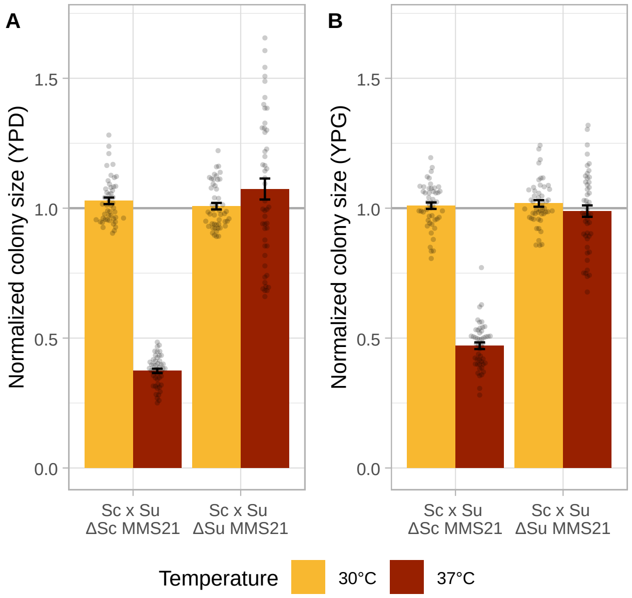


Figure S7: Normalized mean colony sizes of *MMS21* reciprocal hemizygotes on dextrose (A) and glycerol (B) measured after three days of growth at 30 and 37 °C. Whiskers represent standard error of the mean.

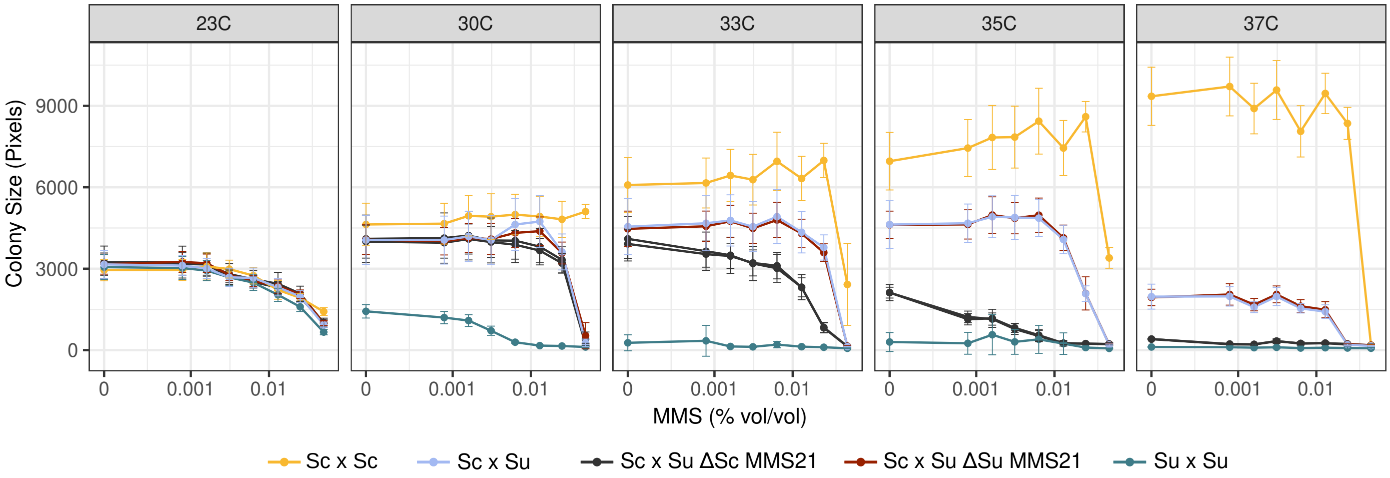


Figure S8: *MMS21* reciprocal hemizygotes' sensitivity to temperature and methyl methanesulfonate (MMS). Colony size of reciprocal hemizygotes are shown relative to *S. cerevisiae* diploid and *S. uvarum* diploid at varying concentrations of MMS after two days of growth. Two independent transformants are shown for ΔSc MMS21.

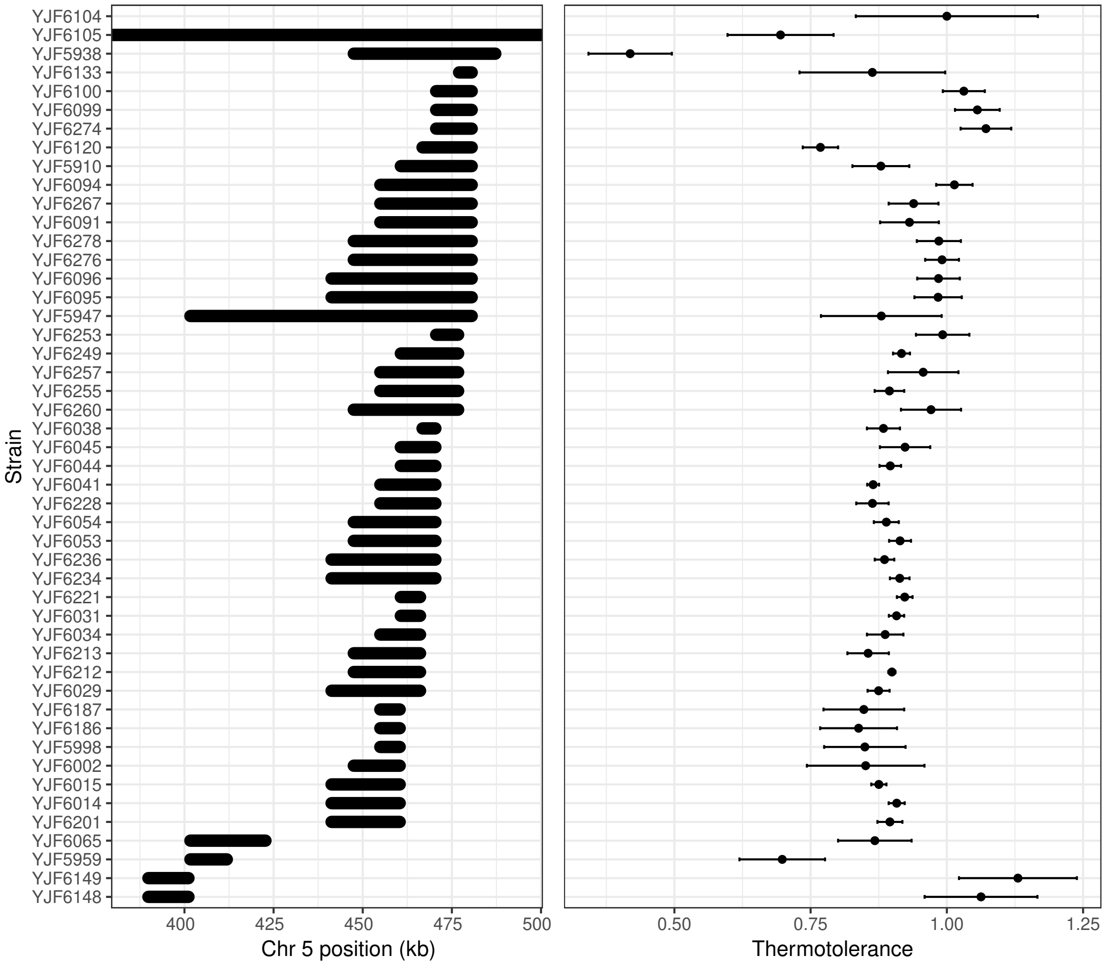


Figure S9: Thermotolerance of *S. cerevisiae* Chr 5 deletions. (A) Chr 5 regions eliminated from the *S. cerevisiae* chromosome in each hybrid strain. Chromosome coordinates are from YPS163 (CASBMI010000006.1). YJF6105 lacks the right arm of Chr 5 and YJF6104 is the euploid control. *BEM2* is located at 480.7-487.2 kb. (B) Thermotolerance was measured by colony size at 37 °C relative to 30 °C after three days of growth on YPD and then scaled to the euploid strain (YJF6104). Error bars show standard deviation.

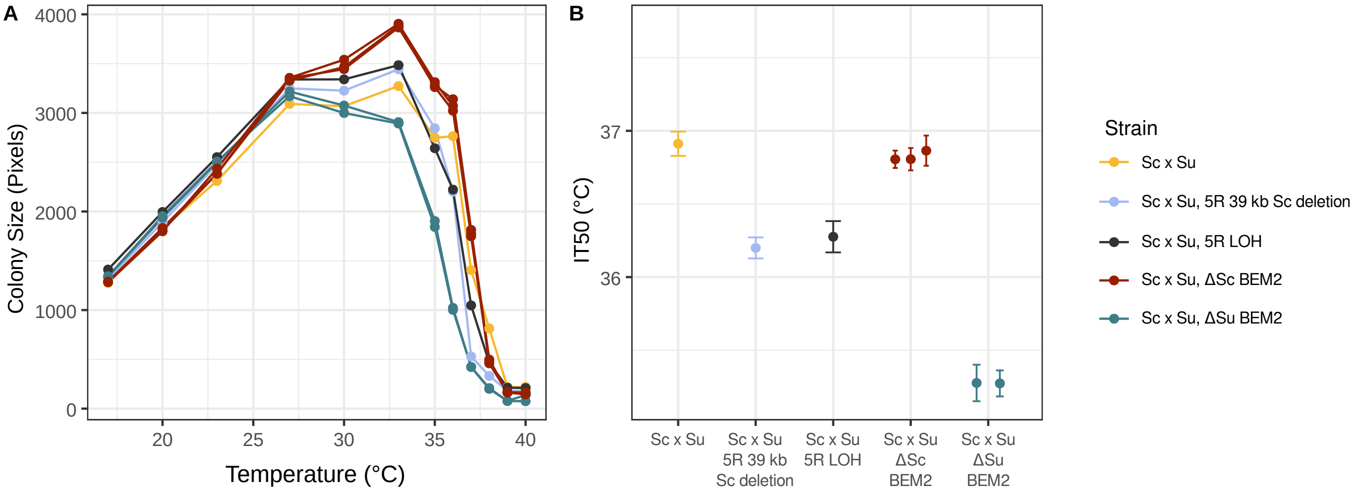


Figure S10: Thermotolerance of *BEM2* reciprocal hemizygotes. Thermotolerance was measured by colony size after five days growth at different temperatures. (A) Mean colony size. (B) Temperature at which growth was inhibited by 50% (IT50). Points and error bars show strain means and standard deviation from a logistic regression. Strains with the 39 kb deletion used to identify *BEM2* (YJF5938) and the loss of *S. cerevisiae* Chr 5R (GS7-D2) are shown for reference. All three replicates of Sc x Su, ΔSc *BEM2* were aneuploid due to elevated coverage of *S. cerevisiae* Chr 1, 9 and 14. The replicate strains of Δ*BEM2* were generated from independent mating events of the same parental strains.
